# Notes on the threatened lowland forests of Mt Cameroon and their endemics including *Drypetes njonji* sp. nov., with a key to species of *Drypetes* sect. Stipulares (Putranjivaceae)

**DOI:** 10.1101/825273

**Authors:** Martin Cheek, Nouhou Ndam, Andrew Budden

**Affiliations:** Royal Botanic Gardens, Kew, Richmond, Surrey, TW9 3AE, UK; Tetra Tech ARD - West Africa Biodiversity & Climate Change (WA BiCC) Program PMB CT58 Accra, Ghana

**Keywords:** Conservation, Cross-Sanaga Interval, South West Region Cameroon, TIPA

## Abstract

**Background and aims:** This paper reports a further discovery of a new endemic threatened species to science in the context of long-term botanical surveys in the lowland coastal forests of Mount Cameroon specifically and generally in the Cross River-Sanaga interval of west-central Africa. These studies focus on species discovery and conservation through the Tropical Important Plant Areas programme.

**Methods:** Normal practices of herbarium taxonomy have been applied to study the material collected. The relevant collections are stored in the Herbarium of the Royal Botanic Gardens, Kew, London and at the Limbe Botanic Garden, Limbe, and the Institute of Research in Agronomic Development – National Herbarium of Cameroon, Yaoundé.

**Key results:** New species to science continue to be discovered from Mt Cameroon. Most of these species are rare, highly localised, and threatened by habitat destruction. These discoveries increase the justification for improved conservation management of surviving habitat. *Drypetes njonji* is described as an additional species new to science and appears to be endemic to the lowland coastal forests at the foot of Mt Cameroon. It is placed in and keyed out in Sect. Stipulares, a group confined to Lower Guinea. The conservation status of *Drypetes njonji* is assessed as Endangered (EN B1+2ab(iii)) according to the 2012 criteria of IUCN. An updated overview of the lowland plant endemics of Mt Cameroon is presented.

## INTRODUCTION

The new species described in this paper as *Drypetes njonji* was brought to light as a result of botanical surveys to aid conservation management at and around Mount Cameroon (Cheek & Hepper, 1994, Cheek *et al.* 1996), long considered a Centre of Plant Diversity (Cheek et al. 1994). The species is a small tree or shrub, an endemic of the coastal forests at the foot of Mt Cameroon. In this paper we provide the evidence that this species is new to science, formally describe it, place it within the current classification of the genus, and review the ecology, endemics, conservation importance and threats to the largely unprotected coastal forest of Mt Cameroon in which it occurs.

### Putranjivaceae and *Drypetes* Vahl

The family Putranjivaceae, formerly included in Euphorbiaceae, is considered to have up to three genera, *Putranjiva* Wall. (four species, restricted to Asia from India to New Guinea), *Sibangea* Oliv. (three species in tropical Africa) and the most species-diverse, *Drypetes* Vahl which is pantropical. *Putranjiva* and *Sibangea* are sometimes subsumed into *Drypetes*. *Drypetes* is considered to have about 200 species, with only c. 20 in the Americas, the remainder in Africa (c. 70spp.) and Asia and Australasia (c. 100 spp. (Hoffman in Heywood *et al.* 2007).

Plants of the World Online (Plants of the World Online, continuously updated) gives 96 accepted names in *Drypetes* for ‘Africa’ but 15 of these are endemic either to Madagascar, Mascarenes, Comores or Seychelles, with 2 in Saõ Tomé, so about 79 taxa are accepted for continental Africa. Only five species occur in S. Africa (e.g. *D. natalensis* (Harv.) Hutch.) while 25 species were accepted by Keay (1958) for Flora of West Tropical Africa. Cameroon with 31 species (if *Sibangea* is included) is the most species diverse African country for *Drypetes* (Onana 2011) followed by Gabon with 26 species (Sosef *et al.* 2006).

*Drypetes* are evergreen, dioecious shrubs and small trees recognised by their alternate, distichous, more or less asymmetric, stipulate, often toothed leaves, and flowers which lack petals, and have 2– 7 unequal, imbricate sepals. The male flowers have 2–20(–50) free stamens with longitudinal dehiscence arranged around a central disc. In the female flower, stamens are absent and the pistil sits on the disc. Carpels are (1–)2(–6), each with 2 axile ovules, which develop into drupaceous fruits (Hoffman in Heywood *et al.* 2007, Radcliffe-Smith 1987:88).

In continental Africa, *Drypetes* species are mainly confined to the understorey of evergreen, usually lowland, forest. They are indicators of good quality, undisturbed forest, in the same way as are species of *Cola* (Cheek pers. obs. 1984–2012, Cheek 2002). Both genera are slow-growing shrubs and trees, not pioneers, and do not regenerate well after major forest disturbance. High species diversity in these genera at a site indicates forest of high conservation value in tropical Africa. Several species in Cameroon have small global ranges and have been assessed as threatened, e.g. *Drypetes magnistipula* Hutch.(EN), *D. preussii* Hutch. (VU), *D. tessmanniana* Pax & K.Hoffm. (VU) (Cheek 2004a, Cheek 2004b, Cheek & Cable 2000) and once revisionary work is completed, the number of threatened species of *Drypetes* is expected to rise (Cheek in Onana & Cheek 2011:144).

Only two new species of *Drypetes* have been described in Africa in the 21^st^ century. These are *D. moliwensis* Cheek & Radcl.-Sm. (Cheek et al. 2000) and *D. batembei* D.J. Harris & Wortley (2006). This low number is partly because of the difficulty caused by the destruction of many German type specimens of *Drypetes* from Central Africa at B in 1943. Loss of this reference material especially hampers the elucidation of new *Drypetes* material from Central Africa because the former German colony of Kamerun was the source of many if not most of these type specimens. It is also here that the genus is most species-diverse, and where discoveries of novelties are therefore still most likely. Currently, newly collected specimens of *Drypetes* often remain unidentified to species. In the Gabon checklist, while 146 specimens are identified to species, a further 147 remained unidentified (Sosef *et al.* 2006).

Burkill (1994:54–60) reports on the uses of 19 species of *Drypetes* in the West African region. The wood of many species is reported as hard and durable, resistant to termites and is valued for constructing homes and mortars for pounding food. The fruits of several species are reported as edible and sometimes sold on markets. The majority of the uses reported are for treating a large variety of ailments, from tooth-ache, wounds, internal and external parasites, fevers, rheumatism, boils and to relieve pain. Some species are used as fish or rat poisons. Johnson et al. (2009) report on the presence of “mustard oil” volatiles in *Drypetes*.

## MATERIAL AND METHODS

The methodology for the surveys in which this species was discovered is recorded in Cheek & Cable (1997). Nomenclatural changes were made according to the Code (Turland et al. 2018). Names of species and authors follow IPNI (continuously updated). Herbarium material was examined with a Leica Wild M8 dissecting binocular microscope fitted with an eyepiece graticule measuring in units of 0.025 mm at maximum magnification. The drawing was made with the same equipment with a Leica 308700 camera lucida attachment. Specimens were inspected from the following herbaria: BM, K, P, WAG, YA. The format of the description follows those in other papers describing new species in *Drypetes* e.g. Cheek et al. (2000). All specimens cited have been seen unless indicated “n.v.”. The conservation assessment follows the IUCN (2012) categories and criteria. GeoCAT was used to calculate red list metrics (Bachman et al. 2011). Herbarium codes follow Index Herbariorum (Thiers, continuously updated).

## RESULTS

The specimens on which the new species, *Drypetes njonji*, is based had been provisionally identified as “*Drypetes ?principum*” in preparation for publication of the Conservation Checklist for Mount Cameroon (Cable & Cheek 1998). But, by oversight, they were not included in that work. Both species have large leaf-blades of similar size and shape and which dry dark brown on the lower surface and bear densely brown hairy fruits of about equal size from the leafy branches. However, the two species can be easily separated using the information in table 1.

**Table. 1.** Characters differentiating between the similar species *Drypetes principum* and *D. njonji*.

|  | *Drypete principum* | *Drypetes njonji* |
| --- | --- | --- |
| Habit | Tree (4–)10–28 m tall | Tree or shrub 2.5–6 m tall |
| Indumentum and colour (dried) of distal stem nodes | Distalmost 1–4 internodes drying matt black, sparsely covered (c. 10–20% of surface) in appressed red hairs | Distalmost internode (below apical bud) completely covered (100% of surface), in golden-brown appressed hairs. Subsequent internodes with white glossy rapidly glabrescent epidermis |
| Leaf-blade dimensions and margins | 12–22 cm long, 5–9 cm wide, margin with numerous teeth | (19.5–)20.5–33(–40.2) cm long, 6.5–11.4(–13) cm wide, margin entire |
| Stipules | Minute, extremely caducous, triangular, <1 mm long. | Conspicuous, persistent up to the fourth node below stem apex, oblong to lanceolate, 9–10 x 3–4 mm, leathery. |
| Male flowers | Stamens 8–12 | Stamens 4 |
| Stigmas in fruit | Flat, obtriangular, 3 mm wide, on styles 1 mm long | Transversely oblong-ellipsoid (sausage-like), sessile, 0.8–0.9 mm long, 1–1.25 mm wide |
| Indumentum of fruit | Patent, woolly appearance | Appressed to surface |

### *Drypetes* Sect. *Stipulares* Pax & K. Hoffm

A remarkable feature of *Drypetes njonji* are the large leathery persistent stipules. In most species of the genus the stipules are vestigial, triangular <1 mm long and early caducous. Persistent stipules are only found in a minority of the species of the genus, most of which occur in West-Central Africa and which in the latest classification of the genus (Pax & Hoffmann 1922) were grouped in Sect. *Stipulares*. This section is characterised by medium to large persistent stipules 5–60 mm long, stamens 4–13 encircling a central disc which lacks a rudimentary ovary. In the female flowers the ovary is 2(–3)–locular, lacks sculpture and is placed on a spreading disc. The stigmas are sessile or subsessile, dilated and undivided (Pax & Hoffmann 1922). *Drypetes njonji* fits this description. A key to the species with persistent stipules is presented below. In this, our new species keys out in a couplet with *D. similis* (Oliv.) Hutch. This species was previously segregated with *D. arborescens* Hutch. as the genus *Sibangea* but was united with *Drypetes* by Hutchinson (1912) who also grouped these persistent stipuled species together as his species 1–7.

Pax & Hoffman (1922) followed Hutchinson in sinking *Sibangea*, but placed *D. similis* in sect. *Hemicyclia* (Wright & Arn.) Pax & K. Hoffm. on account of the unilocular ovary, despite the large, persistent stipules. We have added to this group *Drypetes dinklagei* (Pax) Hutch. since this species also has conspicuous persistent stipules which seem to have been overlooked by Pax & Hoffmann (1922). It is possible that this group is a natural monophyletic unit. Apart from the traits listed by Pax & Hoffmann (1922), these species are all geographically coherent, occurring from eastern Nigeria through Cameroon and Gabon, with some species e.g. *D. mildbraedii* (Pax) Hutch. in DRC. These species also share leaves that completely or mainly lack, the marginal teeth that often characterise the genus, and have leaves that dry brown on the lower surface rather than grey-green. The last feature also occurs in some species outside the group, such as in *D. principum*. Molecular phylogenetic research is needed to test the monophyly of this group.

### Key to the species of Drypetes with large, persistent stipules – *Drypetes* sect. *Stipulares* Pax & K. Hoffm

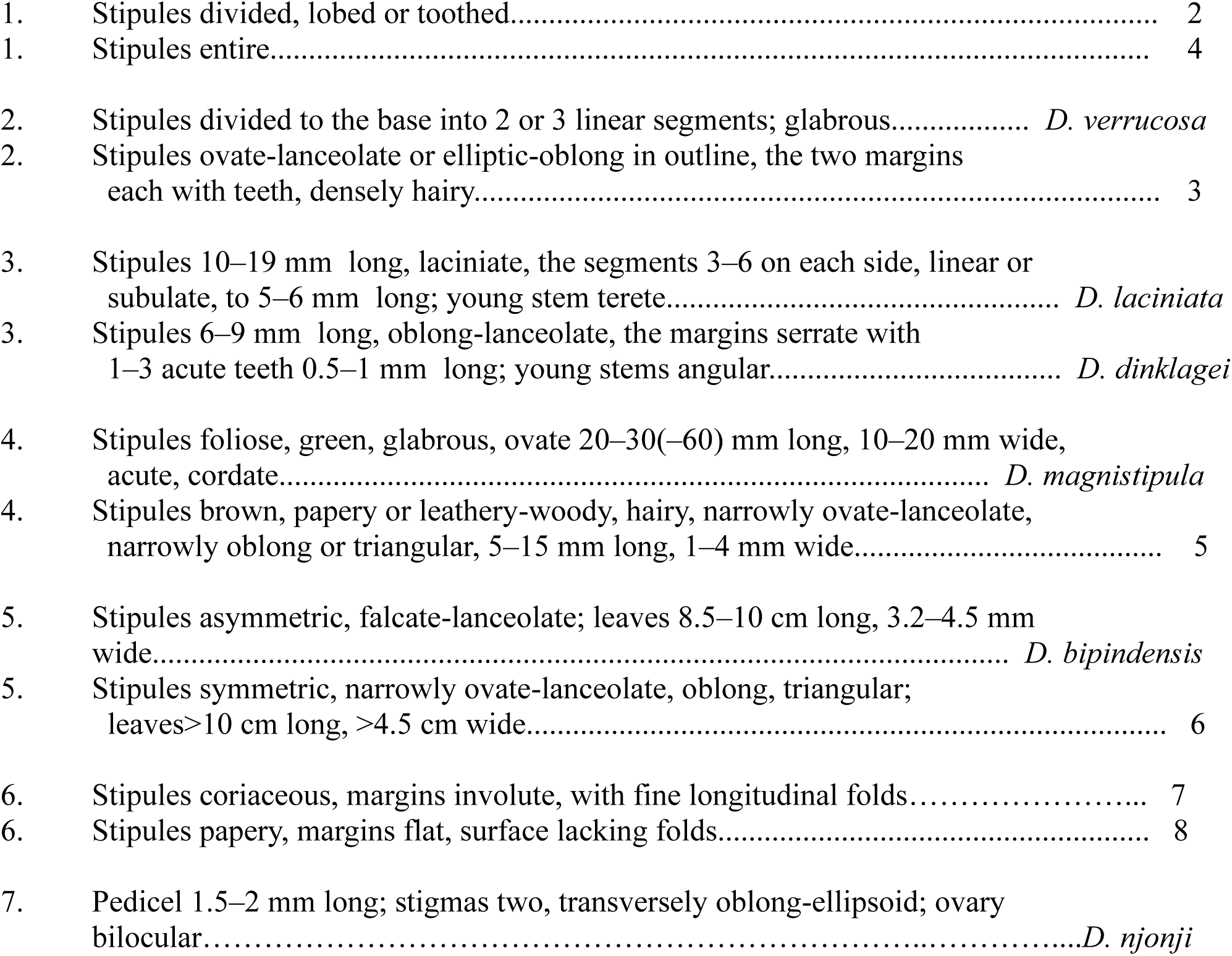
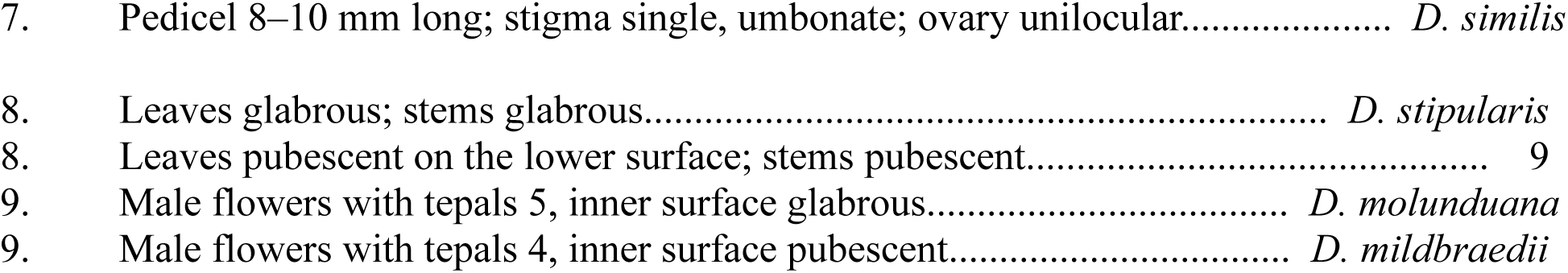

### *Drypetes njonji* Cheek, sp. nov

Differing from *Drypetes principum* Hutch. in the conspicuous, persistent, large (9–10 x 3–4 mm) leathery stipules (not inconspicuous, caducous, small, <1 mm long); distalmost internode completely covered in golden hairs, subsequent nodes glabrous, white (not with all the distal nodes drying black, sparsely covered in red hairs); stigmas sessile, transversely oblong-ellipsoid (not stalked, flat and obtriangular). Type: Cameroon, Southwest Region, on Hunters path to ‘Lake Njonji’, 40 minutes walk N then E from Njonji, 4°08’N, 9°01’E, alt. 150–300 m, fr., 19 Nov. 1993, *Cheek* 5490 (holo–: K barcode K000593141, iso–:BR, K, MO, SCA, WAG, YA).

<u>Dioecious tree or shrub</u> 2.5–6 m tall, trunk 4–7 cm diam. at 1.5 m from the ground, bark grey. Fruiting and flowering from burrs on the leafy stems. Leafy stems distichous, densely golden hairy at completely covering terminal internode of stem apex, rapidly caducous at the internodes below, hairs simple, 0.05–0.2 mm long appressed; epidermis below the first node longitudinally furrowed, grey white or white, glossy, often c. 50% covered in micro-epiphytes, lenticels developing from c. fourth internode, white, slightly raised, elliptic 1–2 mm long, c. 0.5–1 mm wide; internodes even in length 2.1–5.8(–6.3) cm long, diam. 0.3–0.35 cm at the fourth internode from apex, stems slightly flexuose, or terete, with up to 8 leaves per stem. <u>Leaf-blades</u> drying grey-green above, brown below, narrowly oblong-elliptic, rarely slightly oblanceolate-oblong, (19.5–)20.5–33(–40.2) x 6.5–11.4(– 13) cm, acumen 0.4–1.1(–1.4) cm long, acute, base acute, inconspicuously asymmetric, margin entire, lateral nerves (7–)8–11(–12) on each side of the midrib, drying black, arising at 45° from the midrib, arching upwards distally, then running parallel to the margin and connecting with the secondary nerve above by 3–4 brown tertiary nerves, sometimes with a short brown intersecondary nerve present; quaternary nerves brown (concolorous with blade) forming a reticulate pattern, conspicuous on abaxial surface, glabrous. Petiole drying grey, slightly laterally compressed, adaxial surface slightly and shallowly grooved or not, surface longitudinally wrinkled, (0.7–)0.9–1.3 x 0.2 cm, glabrous. <u>Stipules</u> persisting to the fourth node below stem apex, coriaceous, grey-brown, lanceolate or oblong-lanceolate, (7–)8–10 x (2–)3–3.5 mm, apex long tapering, acute, margins revolute, surface finely longitudinally furrowed, indumentum as stem apices. <u>Female Inflorescences</u> in axillary fascicles subtended by the most distal (oldest) leaves from the apex or sometimes at leafless nodes, 1–8-flowered, each flower subtended by a bract and pair of bracteoles. Bracts concave, brown, ovate-elliptic 1.5–1.75 mm long 1–1.25 mm wide, apex acute, completely covered in appressed, yellow-brown hairs, hairs simple, 0.125–0.175 mm long; bracteoles triangular, 0.75–1 mm long and wide, indumentum as bracts. <u>Female flower</u> “with pale yellow ovary and narrow white annular disc” (*Thomas* 9710), 5–7.5 mm long, 3.2–3.5 mm wide, pedicel 1.5–2 mm long, 0.8–1 mm diam., indumentum as bracts. Outer tepals 2, concave, elliptical 3.5–4 mm long, 2.8–3.8 mm wide, apex rounded, outer and inner surface densely covered (90/100% of epidermis covered) in indumentum as bracts but hairs 0.25 mm long. Inner tepals 2, ovate-elliptic, 5.5 mm long, 4.8– 5.5 mm wide, apex and indumentum as outer sepals. Disc protruding beneath ovary by 0.25 mm, 1.25 mm high, rounded, entire, glabrous. Ovary ovoid 5.5–6 mm long, 4.7–6 mm wide, lacking grooves, completely covered in appressed yellow-brown hairs c. 0.25 mm long; 2-locular. Stigmas 2, black, sessile, parallel, transversely oblong-ellipsoid, 0.8–0.9 mm long, 1–1.25 mm wide. <u>Male inflorescences</u> as the female, 1–7-flowered. <u>Male flowers</u> “Calyx light green” (*Tchouto* 687) 4–4.5 mm long, 2.5–3 mm wide at anthesis, opening from globose buds. Pedicel 1.5–2 mm long, 0.5–0.7 mm diam., indumentum as in females. Outer tepals 2, concave, ovate-elliptic, 3.5–4 mm long, 4 Stamens 4, not exserted, 1–1.4 mm long, filaments stout, 0.4 mm long, glabrous, anthers orbicular, 1 mm long, 1 mm wide, dorsal face flattened, ventral with 4 longitudinal thecae, apex rounded, apex with 2–3 tufts of hairs 0.5 mm long, margin fringed with hairs. Disc flat, glabrous, rounded-quadrangular 1.75 mm diam., each of the 4 edges with a shallow notch (accommodating an anther filament), upper surface with raised coarse reticulations, cells c. 0.2–0.3 mm diam. Rudimentary ovary absent. <u>Fruit</u> ellipsoid, (1.2–)1.4–1.9 cm long, 0.8–0.9(–1) cm diam., with 2 opposite shallow longitudinal grooves when dry, completely covered in appressed whitish brown hairs. Stigmas persistent, not accrescent. Pedicel accrescent (0.15–)0.2–0.4 cm long. Tepals persistent, not accrescent. <u>Seeds</u> 2, one per locule, dark brown, plano-convex, 16 mm long, 10 mm wide, 6 mm broad, lacking surface ornamentation. Fig. 1, 2.

**Figure 1.**
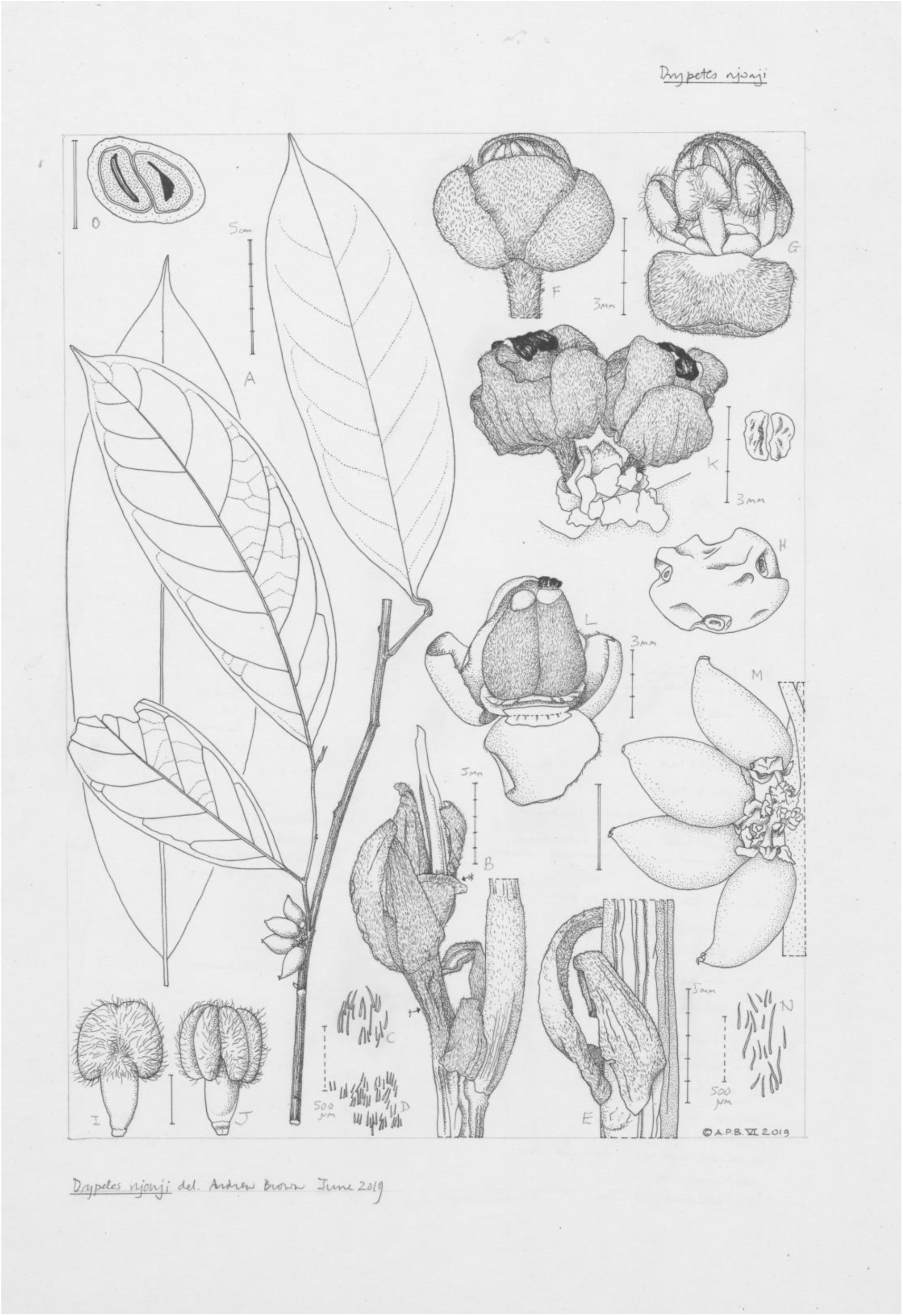
*Drypetes njonji*. **A.** Habit including fruiting shoot. Outline of larger leaf in background. **B.** Stem apex. **C.** Detail of hairs at *. **D.** Detail of hairs at Ⴕ. **E.** Pair of stipules. **F.** Male flower, side view (hydrated). **G.** Male flower in F with outer sepals removed and front inner tepal reflexed to show stamens. **H.** Disc from F viewed from above. **I.** Stamen, outer face. **J.** Stamen, inner face. **K.** Female flowers *in situ* (dry) with view of stigmas from above. **L.** Side view of young fruit *in situ* (hydrated) [hairs on inner and outer faces of sepals omitted]. **M.** Fruits *in situ*. **N.** Hairs from fruit surface. **O.** Mature fruit, TS (hydrated). A-E, K, M-O from *Cheek et al. 5490* (K). F-J from *Tchouto 687* (K). L from *Ndam* 754 (K). Scale bars: A = 5 cm; B & E = 5 mm; C-D & N = 500µm; F-H & K = 3 mm; I-J = 1 mm; M & O = 1 cm. Drawn by ANDREW BROWN.mm wide, apex rounded, indumentum as females covering 20–70% of surface. Inner sepals 2, orbicular 3 mm long, 3 mm wide, indumentum as in females, 90% cover.

#### Habitat and distribution

Cameroon, so far only known from coastal forest of Mt Cameroon in South West Region. Infrequent small tree or shrub of undisturbed lowland evergreen forest on old volcanic or pre-cambrian rocky soils; 50–300m alt.

**Figure 2.**
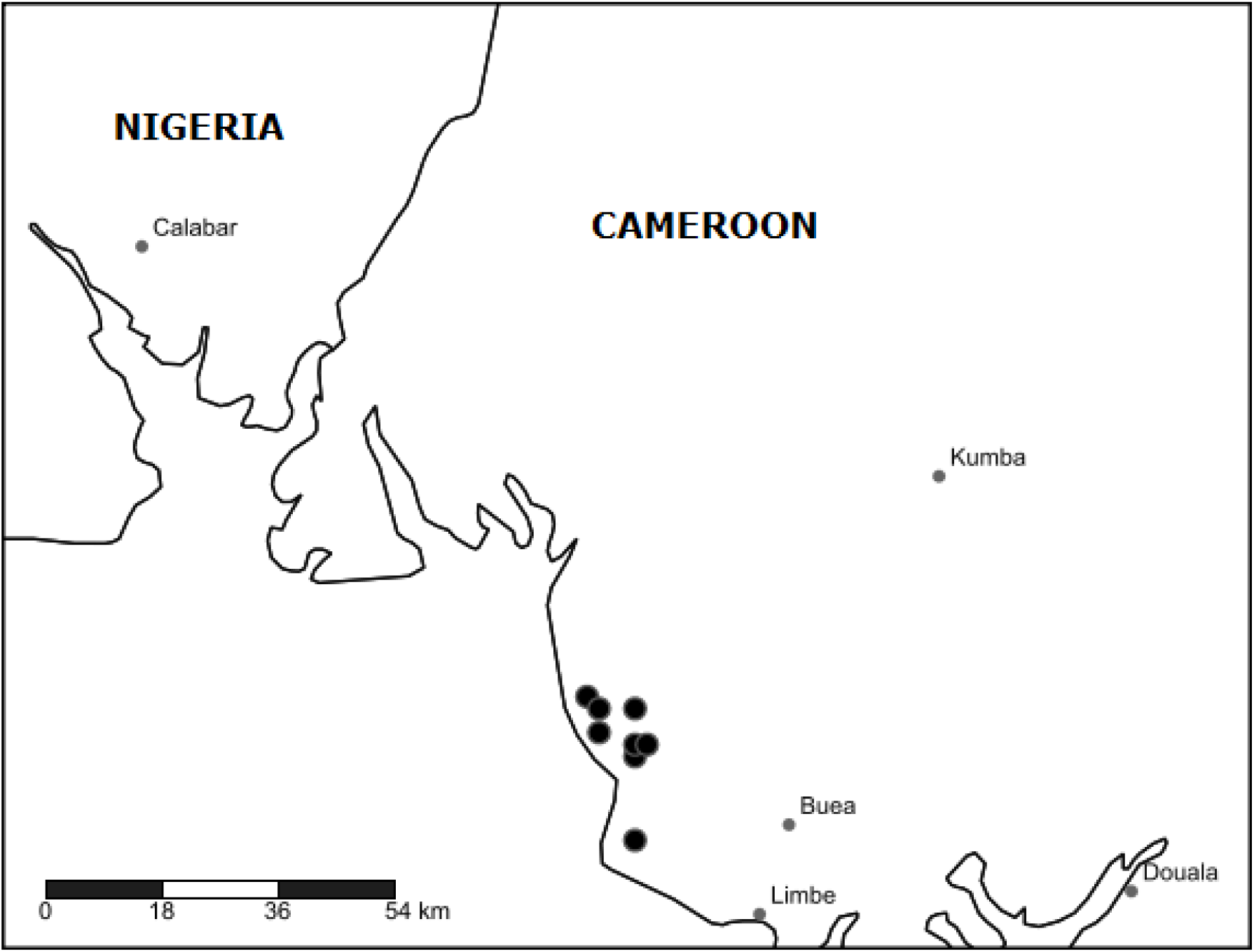
Distribution of *Drypetes njonji*.

#### Additional specimens studied

**Cameroon**: South West Region (formerly Province), **Mount Cameroon, Onge Forest**, Bomana-Koto road c 500 m, bearing 305° towards Onge river, 5 hr walk from the road, 4°19’N, 9°01’E, 400 m alt., fl., fr., 20 Oct 1993, *Ndam* 754 (K, SCA n.v., YA n.v.); Bomana-Koto road c 500 m, bearing 305° towards Onge river, 3 hr walk from the road, 4°19’N, 9°01’E, 400 m alt., fl., fr., 18 Oct 1993, *Ndam* 681 (K, SCA n.v.); Southwest Region, Bomana secondary forest, 04°15’N, 09°01’E, 200 m alt, fr., 5 Oct 1993, *Tchouto Mbatchou* 687, (K, SCA n.v., YA n.v.); Bomana secondary forest, 4°15’N, 9°1’E, 200 m alt, 6 Oct 1993, *Tchouto Mbatchou* 706 (K, SCA n.v.); Onge River secondary forest, 4°16’N, 9°1’E, 240 m alt, 9 Oct 1993, *Tchouto Mbatchou* 706 (K, SCA n.v.); Southwest Region, 0.5 km west of the Idenao-Bomana road, in a ravine formed by the Lokange River, 4°16’N, 9°02’E, 100 m alt., fl., fr., 6 Oct 1993, *Thomas* 9710 (K, SCA n.v., YA n.v.); West bank of the Onge River, 4°17’N, 8°58’E, 100 m alt., 7 Nov 1993, *Thomas* 9809 (K, SCA n.v.); Forested hillside west of the Onge River and ridges on “Thump Mount”, 4°20’N, 8°57’E, 200 m alt., fr., 9 Nov 1993, *Thomas* 9868 (K, SCA n.v.); Forest to the west of Onge River, c. 6 km along the Onge River from its mouth and c. 8 km northwest of Idenau, 4°19’N, 8°58’E, 100 m alt., 11 Nov 1993, *Watts* 1029 (K, SCA n.v.).

#### Etymology

named (noun in apposition) for the community of Njonji from the forests of which the type specimen was collected.

#### Conservation

*Drypetes njonji* has an extent of occurrence of 94 km^2^ and area of occupancy of 32 km^2^ as calculated using GeoCAT (Bachman et al. 2011). There are only two “threat-based” locations (IUCN 2012), in the Onge forest and Bomana-Njonji forest. These areas of Mount Cameroon are threatened with forest clearance, both for timber and oil palm. Species within the genus *Drypetes* are slow growing trees and shrubs and are indicators of good quality, undisturbed forest. They do not regenerate well after major forest disturbance and are therefore especially sensitive to the threats posed by forest clearance. Given the threats and restricted range, an assessment of Endangered (EN B1+2ab(iii)) is given to this species.

#### Notes

*Drypetes njonji* may yet be discovered to have a wider range than Mt Cameroon. Several other species previously considered endemic to the mountain have since been found in other locations (see discussion on endemics of Mt Cameroon below). However, following the intensive survey at Mt Cameroon (Cable & Cheek 1998) during which the specimens cited above of *Drypetes njonji* were collected, surveys of other areas in Cameroon followed, for example at Korup National Park (Cheek & Cable 1998), Mt Kupe and the Bakossi Mts (Cheek et al. 2004), in the Lebialem Highlands (Harvey et al. 2010), the Bali Ngemba Forest Reserve (Harvey et al. 2004) at Dom (Cheek et al. 2010) and at Ebo (Cheek et al. 2018a). Yet no further sites for *Drypetes njonji* were found.

### Mt Cameroon, its endemic plants and its coastal forests

At 4095 m high, Mt Cameroon, locally known as Fako, is by far the highest mountain, and the only active volcano, in continental West-Central Africa. One has to travel about 1300 km eastwards to the Virunga Mts of eastern DRC before encountering mountains of comparable height, or active volcanoes. Fako erupts roughly every 20 years and the last lava flow dates from May 2000.

Rainfall varies dramatically from one part of the mountain to another, most falling in the 8–9 month wet season between April to November. There are no months with less than 50 mm precipitation at Cape Debundscha, on the SW, seaward side, which receives 10–15 m p.a., but at Tiko to the E there are two months with <50 mm, and rainfall declines to 2–3 m p.a. while at the summit and in the rainshadow to the E, it is 1–2 m p.a (Courade 1974).

The volcanic massif is about 45 km along its longest axis, SSW-NNE, and 28 km across at its narrowest (Cable & Cheek 1998). Together with the foothill areas and lower slopes that surround it, it has the highest documented species diversity for vascular plants in tropical Africa, with 2435 species, of which 49 have been considered strict endemics and of which 116 have been considered threatened (Cable & Cheek 1998). However, numerous additional new species, several endemic, have been published in the last 20 years.

Botanical exploration of Mt Cameroon began in 1861 with the Kew botanist Gustav Mann, resulting in the remarkable paper on the high-altitude Flora of the mountain (Hooker 1864). Numerous other botanists including Preuss, Schlechter, Mildbraed, Kalbreyer, Dusen, Maitland, Brenan and Letouzey followed Mann in increasing knowledge of the plants of this mountain (Cable & Cheek 1998).

The Mount Cameroon National Park was created in 2009, to a large degree because of its unique botanical importance. The mountain is a tourist attraction, particularly for those who wish to walk to the summit from the former German colonial capital of Buea on the eastern slope of the massif. Forest elephants and the endemic Mt Cameroon Francolin (*Pternistus camerunensis*) are also attractions, although rarely seen.

Arguably the most spectacular botanical discovery at Mount Cameroon, from its northern foothills, the S.Bakundu Reserve, was that of *Medusandra richardsiana* Brenan, basis of a new family, the Medusandraceae (Brenan 1952), to which was later added the genus *Soyauxia* Oliv. *Medusandraceae* sensu stricto (*Medusandra* Brenan) was until recently considered endemic to the Cross-Sanaga interval (Heywood et al. 2007). Subsequently, however, both *Soyauxia* and later *Medusandra* were shown to be confamilal with the rare and little known S.American family, Peridiscaceae (Soltis et al. 2007, Breteler et al. 2015). The Cross-Sanaga interval (Cheek et al. 2001), comprising largely of South West Region, Cameroon, contains the area with the highest species diversity per degree square in tropical Africa (Barthlott et al. 1996, Cheek et al. 2006), and many of these species are narrow endemics, and a large number are at Mt Cameroon.

The emphasis of the UK government Mt Cameroon Plant Genetic Resources Project, later known as the Mount Cameroon Project was to focus on conservation management of the forest plant diversity, especially the highly threatened lowland forest vegetation. Following rehabilitation of the Limbe Botanic Garden, its base, the project conducted surveys in the eastern foothills formerly known as Mabeta-Moliwe (Cheek 1992) now known as Bimbia-Bonadikombe, and then of the Etinde area (Thomas and Cheek 1992), and in the coastal forests between Idenau and Limbe, with the adjoining coastal eastern foothill forest of Onge (late 1993) and finally Mokoko forest (early 1994). Following identifications, a checklist for Mt Cameroon and its foothills was published (Cable & Cheek 1998). The specimens of *Drypetes njoni* cited in this paper were collected during these surveys.

Several Mount Cameroon endemics occur above 2000 m alt. both at the forest-grassland ecotone or in the montane grassland above that begins the summit area. These include *Silene biafrae* Hook.f. (Hooker 1864), *Myosotis cameroonensis* Cheek & R. Becker (Cheek & Becker 2004) and *Luzula mannii* (Buchenau) Kirschner & Cheek subsp. *mannii* (Kirschner & Cheek 2000). Yet some species among the nine previously listed as endemic to this summit area (Cable & Cheek 1998) are now known to have a wider distribution. Both *Genyorchis macrantha* Summerh. and also *Isoglossa nervosa* C.B.Cl. have subsequently been found on another peak in the Cameroon Highlands, Mt Oku (Darbyshire et al. 2011), which at 3095 m alt, is also a centre of plant diversity of conservation importance (Maisels et al. 2000, Cheek et al. 2000). Yet a greater number of the Mt Cameroon endemics are found in the submontane or cloud forest that extend from c. 800–2000 m altitude (Tchouto *et al.*1999; Thomas & Cheek 1992). Eleven are listed in Cable & Cheek (1998), several of which were only formally named and published subsequently. These include *Oxygyne duncanii* Cheek (Cheek *et al.* 2018a) *Impatiens etindensis* Cheek & Eb. Fisch (Cheek & Fischer 1999), *Impatiens frithii* Cheek (Cheek & Csiba 2002a) and orchids such as *Angraecopsis cryptantha* P.J. Cribb (Cribb 1996). However, most of the Mt Cameroon endemics and the most threatened species overall, are found in the evergreen lowland forest that survives in the foothills and in coastal areas below 800 m altitude. Twenty-nine species of lowland endemics are listed by Cable & Cheek (1998: xxxii) of which 17 had been newly discovered and still awaited publication. Subsequently, some of these supposed endemic species were either synonymised, e.g. *Trichoscypha camerunensis* Engl. or found to have wider distributions, e.g. *Trichoscypha bijuga* Engl. (Breteler 2004). Some of those subsequently published as new to science and thought to be endemic, such as *Salacia nigra* Cheek (Gosline & Cheek 2014) and *Belonophora ongensis* S.E.Dawson & Cheek, were later found to extend beyond Mt Cameroon to Littoral Region (Cheek et al. 2018b, Cheek & Dawson 2000). This was also the case with *Stelechantha arcuata* S.E. Dawson (2002) now *Pauridiantha arcuata* (S.E. Dawson) Smedmark & B. Bremer (2011) extending to Mt Kupe and the Bakossi Mts (Cheek et al. 2004), *Psychotria elephantina* Lachenaud & Cheek (Cheek & Lachenaud 2013) extending to Korup (Lachenaud 2019), and also *Ancistrocladus grandiflorus* Cheek (Cheek 2000), subsequently found to extend to the Rumpi Hills.

But other endemics recorded for Mt Cameroon in Cable & Cheek (1998), under working names that were formalised later, such as *Cola cecidifolia* Cheek (Cheek 2002), *Psychotria bimbiensis* Bridson & Cheek (Cheek & Bridson 2002), *Drypetes moliwensis* Cheek & Radcl.-Sm. (Cheek et al. 2000) remain strictly endemic to the coastal lowland forests of Mt Cameroon. Re-examination of specimens collected in the surveys of the 1990s has uncovered further endemic new species, not listed as such in Cable & Cheek (1998), such as *Dracaena mokoko* Mwachala & Cheek (Mwachala & Cheek 2012) and *Octoknema mokoko* Gosline & Malécot (Gosline & Malécot 2012). New surveys at Mt Cameroon in the 21^st^ century have discovered other highly threatened species entirely overlooked in the 1990s, such as *Kupea martinetugei* Cheek (Cheek et al. 2003), *Psychotria asterogramma* Lachenaud (Lachenaud 2019), and a further new endemic *Afrothismia foertheriana*, T.Franke, Sainge & Agerer (Franke et al. 2004).

The rarest of the rare endemic Mt Cameroon plant species were considered those nine species listed as possibly extinct globally in Cable & Cheek (1998) since they had not been seen for over 60 years (now 80 years), not being found in the surveys of the 1990s. Eight of these species were lowland forest endemics. In one case, *Plectranthus dissitiflorus* (Guerke) J.K. Morton, (now *Coleus dissitiflorus* Guerke) a collection was discovered from the 1970s giving hope that it survives today. But the remainder have not been seen leading to fears that they are indeed extinct. In one case, *Oxygyne triandra* Schltr. a series of concerted efforts were made to rediscover it over several years, without success, leading to the firm conclusion that it is indeed now globally extinct (Cheek et al. 2018a). This is also the case with a further mycotroph, *Afrothismia pachyantha* Schltr., that had been collected with the *Oxygyne.* This *Afrothismia* was thought to have been locally extinct at Mt Cameroon but was considered to have been rediscovered on Mt Kupe (Cheek 2004c), until the Mt Kupe population was found to be a separate, species rendering *Afrothismia pachyantha* globally extinct (Cheek et al. 2019).

The coastal forest habitats of all these coastal lowland forest species fall outside the Mount Cameroon National Park which is restricted to higher altitude. Lowland forest and its species are undoubtedly the most threatened at Mt Cameroon. Clearance of coastal forest had begun in the eastern foothills area of Mabeta by the early 19^th^ century to produce food for slaves being held at Bimbia by the Isuwu. Later the Christian settlement of Victoria (now Limbe) was founded in 1858 at Ambas Bay by Alfred Saker of the London Missionary Society. More extensive forest clearance began with the advent of the German colony of Kamerun in 1884. The fertile volcanic soils and abundant rainfall merited extensive lowland forest clearance to support plantation crops for export to Europe, initially of bananas, later of rubber and oil palm. Oil palm plantation now extends in a belt around the seaward southern base of Mount Cameroon from Cameroon’s largest port, Douala, but does not yet occupy the more infertile western foothill areas. Today, the lowland forest areas that remain are threatened by clearance for timber, followed by subsistence agriculture and plantation expansion. Limbe is now a city, and a port, with an oil refinery and a population of about 118,000 persons (Cameroon data portal 2015).

These threats make it a priority to name Mt Cameroon coastal forest endemic species such as *Drypetes njonji*, since until this is done, conservation assessments are not accepted by IUCN (IUCN 2012), and as a result, proposals for conservation measures are less likely to be enacted.

The number of flowering plant species known to science is disputed (Nic Lughadha et al. 2017), but of the estimated 369 000 vascular plant species known (Nic Lughadha et al. 2016), only 7.2% are included on the Red List using the IUCN (2012) standard (Bachman et al. 2019). About 2000 new species are still discovered each year (Willis 2017), and a high proportion of these are likely to be threatened, since more widespread species tend already to have been discovered, although there are exceptions (Cheek & Etuge 2009). This makes it all the more urgent to find, document and protect undescribed species before they become globally extinct, as has *Oxygyne triandra* Scltr. (Cheek et al. 2018a)

Efforts are now being made to delimit the highest priority areas in Cameroon for plant conservation through the Tropical Important Plant Areas (TIPAs): Cameroon programme (Cheek, continuously updated) which uses the revised Important Plant Areas (IPA) criteria set out in Darbyshire et al. (2017). The coastal forests of Mt Cameroon, the habitat for *Drypetes njonji* will be among the TIPAs so designated.

## ACKNOWLEDGEMENTS

Janis Shillito typed the manuscript. George Gosline and two anonymous reviewers gave advice on an earlier version of the manuscript. Fieldwork funding in the 1990s leading to the discovery and collection of most of the specimens cited in this paper was received from the Overseas Development Administration (ODA) of the UK government (now the Department for International Development) through the former Mount Cameroon Project at Limbe Botanic Garden, co-managed by ODA with the Forest Department of the Cameroon Government. The fieldwork during which the type collection was collected was supported by the Earthwatch Institute.

## REFERENCES

Bachman S., Moat J., Hill A.W., de la Torre J., Scott B. (2011) Supporting Red List threat assessments with GeoCAT: geospatial conservation assessment tool, in: Smith V, Penev, eds. e-Infrastructures for data publishing in biodiversity science. ZooKeys 150: 117–126. Available from: http://geocat.kew.org/ [accessed 19 July 2019].

Bachman S.P., Field R., Reader T., Raimondo D., Donaldson J., Schatz G.E., Nic Lughadha E.M. (2019) Progress, challenges and opportunities for Red Listing. Biological Conservation 234: 45–55.

Barthlott W., Lauer W., Placke A. (1996) Global distribution of species diversity in vascular plants: towards a world map of phytodiversity. Erkunde 50: 317–328 (with supplement and figure).

Brenan J. P. M. (1952) Plants of the Cambridge Expedition, 1947–1948: II. A New Order of Flowering Plants from the British Cameroons Kew Bulletin 7(2): 227–236. https://doi.org/10.2307/4109266

Breteler F.J. (2004) The genus Trichoscypha (Anacardiaceae) in Lower Guinea and Congolia: A synoptic revision. Adansonia 26(1) 97–127

Burkill H.N. (1994). The Useful Plants of West Tropical Africa. Vol. 2, families E-I. – Kew, Royal Botanic Gardens.

Breteler F.J, Bakker F.T, Jongkind C.C.H. (2015) A synopsis of Soyauxia (Peridiscaceae, formerly Medusandraceae) with a new species from Liberia. Plant Ecology and Evolution 148: 409–419. https://doi.org/10.5091/plecevo.2015.1040

Burkill H.N. (1994) The Useful Plants of West Tropical Africa. Vol. 2, families E-I. – Kew, Royal Botanic Gardens. https://doi.org/10.2307/4111051

Cable S., Cheek M. (1998) The plants of Mt Cameroon, a conservation checklist. Kew, Royal Botanic Gardens.

Cameroon data portal (2015) http://cameroon.opendataforafrica.org/PHCC2015/population-and-housing-census-of-cameroon-2015?tsId=1002400 (accessed Oct. 2019).

Cheek M. (continuously updated) The Tropical Important Plant Areas (TIPAs): Cameroon programme.https://www.kew.org/science/our-science/projects/tropical-important-plant-areas-cameroon. Downloaded on 06 October 2019.

Cheek M. (1992) A Botanical Inventory of the Mabeta-Moliwe Forest. Report to Govt. Cameroon from O.D.A. Kew, Royal Botanic Gardens.

Cheek M. (2000) A synoptic revision of Ancistrocladus (Ancistrocladaceae) in Africa, with a new species from western Cameroon. Kew Bulletin 55: 871–882. https://doi.org/10.2307/4113632

Cheek M. (2002) Three new species of Cola (Sterculiaceae) from western Cameroon, Cameroon. Kew Bull. 57: 402–415. https://doi.org/10.2307/4111117

Cheek M. (2004a) Drypetes preussii. The IUCN Red List of Threatened Species 2004: e.T34775A9888700. http://dx.doi.org/10.2305/IUCN.UK.2004.RLTS.T34775A9888700.en. Downloaded on 04 October 2019.

Cheek M. (2004b) Drypetes magnistipula. The IUCN Red List of Threatened Species 2004: e.T45451A10999341. http://dx.doi.org/10.2305/IUCN.UK.2004.RLTS.T45451A10999341.en. Downloaded on 04 October 2019.

Cheek, M. (2004c) Afrothismia pachyantha. The IUCN Red List of Threatened Species 2004: e.T39539A10246294. http://dx.doi.org/10.2305/IUCN.UK.2004.RLTS.T39539A10246294.en. Downloaded on 06 October 2019.

Cheek M., Becker R. (2004) A new species of Myosotis L. (Boraginaceae) from Cameroon, with a key to the Tropical African species of the genus. Kew Bulletin 59: 227–231. https://doi.org/10.2307/4115854

Cheek M., Bridson D. (2002) Two new species of Psychotria (Rubiaceae) from western Cameroon. Kew Bulletin 57: 389–395. https://doi.org/10.2307/4111114

Cheek M., Cable S. (1997) Plant Inventory for conservation management: the Kew-Earthwatch programme in Western Cameroon, 1993-96. In: Doolan S. (Ed.) African rainforests and the conservation of biodiversity: 29–38. Oxford, Earthwatch Europe.

Cheek M., Cable S. (1998). Preliminary results of the botanical inventory of the Ekundu Kundu Region of the Korup Park, pp. 72–80 in Songwe, N. C. (Ed.) Proceedings of Workshop on Korup National Park & Project Area. Mundemba, Cameroon, Korup Project.

Cheek M., Cable S. (2000) Drypetes tessmanniana. The IUCN Red List of Threatened Species 2000:e.T39517A10243683. http://dx.doi.org/10.2305/IUCN.UK.2000.RLTS.T39517A10243683.en. Downloaded on 04 October 2019.

Cheek M., Csiba L. (2002a) A new epiphytic species of Impatiens (Balsaminaceae) from western Cameroon. Kew Bulletin 57(3): 669–674. https://doi.org/10.2307/4110997

Cheek M., Csiba L. (2002b) A revision of the Psychotria chalconeura complex (Rubiaceae) in Guineo-Congolian Africa. Kew Bulletin 57: 375–387. https://doi.org/10.2307/4111113

Cheek M., Dawson S. (2000) A synoptic revision of *Belonophora* Hook. f. (Rubiaceae). Kew Bull. 55: 63–80. https://doi.org/10.2307/4117761

Cheek M., Etuge M. (2009) A new submontane species of Deinbollia (Sapindaceae) from Western Cameroon and adjoining Nigeria. Kew Bulletin 64: 503–508. https://doi.org/10.1007/s12225-009-9132-4

Cheek M., Fischer E. (1999) A tuberous and epiphytic new species of Impatiens (Balsaminaceae) from Southwest Cameroon. Kew Bulletin 54: 471–475. https://doi.org/10.2307/4115828

Cheek M., Hepper F.N. (1994) Progress on the Mount Cameroon Rainforest Genetic Resources Project, in Wildlife Conservation in West Africa. Nigerian Field Society (UK) Occasional Paper No. 1: 15–19

Cheek M., Lachenaud O. (2013) Psychotria elephantina sp. nov. (Rubiaceae) an Endangered rainforest shrub from Cameroon. Nordic Journal Botany 31(5): 569–573. http://dx.doi.org/10.1111/j.1756-1051.2012.01488.x

Cheek M., Achoundong G., Onana J-M., Pollard B., Gosline G., Moat J., Harvey Y.B. (2006) Conservation of the Plant Diversity of Western Cameroon. In: S.A. Ghazanfar, H.J. Beentje (eds) Taxonomy and ecology of African plants, their conservation and sustainable use. Proceedings of the 17th AETFAT Congress, Addis Ababa, Ethiopia: 779–791. Kew, Royal Botanic Gardens.

Cheek M., Cable S., Hepper F.N., Ndam N., Watts J. (1996) Mapping plant biodiversity on Mt. Cameroon. pp. 110–120 in van der Maesen, van der Burgt & van Medenbach de Rooy (Eds), The Biodiversity of African Plants (Proceedings XIV AETFAT Congress). Kluwer. https://doi.org/10.1007/978-94-009-0285-5_16

Cheek M., Etuge M., Williams S. (2019) Afrothismia kupensis sp. nov. (Thismiaceae), Critically Endangered, with observations on its pollination and notes on the endemics of Mt Kupe, Cameroon. Blumea - Biodiversity, Evolution and Biogeography Of Plants. 64(1): 158–164 https://doi:10.3767/blumea.2019.64.02.06

Cheek M., Harvey Y., Onana J-M. (2010) The Plants of Dom, Bamenda Highlands, Cameroon, A Conservation Checklist. Kew, Royal Botanic Gardens.

Cheek M., Mackinder B. Gosline G., Onana J., Achoundong G. (2001) The phytogeography and flora of western Cameroon and the Cross River-Sanaga River interval. Systematics and Geography of Plants 71: 1097–1100. https://doi.org/10.2307/3668742

Cheek M., Onana J-M., Pollard B.J. (2000b) The Plants of Mount Oku and the Ijim Ridge, Cameroon, a Conservation Checklist. Kew, Royal Botanic Gardens.

Cheek M., Pollard B.J., Darbyshire I., Onana J-M., Wild C. (2004) The Plants of Kupe, Mwanenguba and the Bakossi Mountains, Cameroon: A Conservation Checklist. Kew, Royal Botanic Gardens.

Cheek M, Prenner G, Tchiengué B, Faden R.B. (2018b) Notes on the endemic plant species of the Ebo Forest, Cameroon, and the new, Critically Endangered, Palisota ebo (Commelinaceae). Plant Ecology & Evolution 151(3): 434–441. https://doi.org/10.5091/plecevo.2018.1503

Cheek M., Radcliffe-Smith A., Faruk A. (2000) A new species of Drypetes (Euphorbiaceae) from Western Cameroon. Kew Bull. 55: 871–882. https://doi.org/10.2307/4113635

Cheek M., Thomas D., Besong J.B., Gartlan S., Hepper F.N. (1994) Mount Cameroon, Cameroon pp. 163–166 in S. D. Davies, V.H. Heywood & A.C. Hamilton (eds.) Centres of Plant Diversity: A Guide and Strategy for their Conservation. Cambridge, WWF/IUCN.

Cheek M., Tsukaya H., Rudall P.J., Suetsugu K. (2018a) Taxonomic monograph of *Oxygyne* (Thismiaceae), rare achlorophyllous mycoheterotrophs with strongly disjunct distribution. PeerJ 6: e4828. https://doi.org/10.7717/peerj.4828

Cheek M., Williams S., Etuge M. (2003) Kupea martinetugei, a new genus and species of Triuridaceae from western Cameroon. Kew Bulletin 58: 225–228. https://doi.org/10.2307/4119366

Courade G. (1974) Commentaire des Cartes. Atlas Regional. Ouest 1. ORSTOM, Yaoundé.

Cribb P.J. (1996) New species and records of orchids from West Africa. Kew Bulletin 51: 353–364. https://doi.org/10.2307/4119329

Darbyshire I., Anderson S., Asatryan A., et al. (2017) Important Plant Areas: revised selection criteria for a global approach to plant conservation. Biodiversity Conservation 26: 1767–1800. https://doi.org/10.1007/s10531-017-1336-6

Darbyshire I., Pearce L., Banks H. (2011) The genus Isoglossa (Acanthaceae) in west Africa. Kew Bulletin 66: 425–439. https://doi.org/10.1007/s12225-011-9292-x

Dawson S. (2002). A New Species of Stelechantha Bremek. (Rubiaceae, Urophylleae) from Cameroon. Kew Bulletin 57: 397–402. https://doi.org/10.2307/4111116

Franke T, Sainge M, Agerer R. (2004) A new species of Afrothismia (Burmanniaceae, tribe Thismieae) from the western foothills of Mt Cameroon. Blumea 49: 451–456. https://doi.org/10.3767/000651904x484397

Gosline G., Cheek M., Kami T.(2014) Two new African species of Salacia (Salacioideae, Celastraceae). Blumea 59: 26–32. https://doi.org/10.3767/000651914x682026

Gosline G., Malécot V. (2012) A monograph of Octoknema (Octoknemaceae — Olacaceae s.l.). Kew Bulletin 66: 367–404. https://doi.org/10.1007/s12225-011-9293-9

Harris D.J., Wortley A.H. (2006) A new species of Drypetes (Putranjivaceae) from the Central African Republic. Edinburgh Journal of Botany 62: 253–256. https://doi.org/10.1017/s096042860600059x

Harvey Y., Pollard B.J., Darbyshire I., Onana J-M., Cheek M. (2004) The Plants of Bali Ngemba Forest Reserve, Cameroon. A Conservation Checklist. Kew, Royal Botanic Gardens.

Harvey Y.H., Tchiengue B., Cheek M. (2010) The plants of the Lebialem Highlands, a conservation checklist. Kew, Royal Botanic Gardens.

Heywood V.H.H. (2007) Medusandraceae. In: Heywood VH, Brummitt RK, Culham A, Seberg O (eds), Flowering plant families of the world: 205. Kew, Royal Botanic Gardens.

Hoffmann P. (2007) Putranjivaceae. In: Heywood VH, Brummitt RK, Culham A, Seberg O (eds), Flowering plant families of the world: 270–271. Kew, Royal Botanic Gardens.

Hooker J.D. (1864) On the plants of the temperate regions of Cameroons Mountain and in islands in the bight of Benin collected by Gustav Mann, government botanist. Journal of the Linnaean Society of London (Botany) 7: 171–240. https://doi.org/10.1111/j.1095-8312.1864.tb01067c.x

Hutchinson J. (1912) Drypetes pp. 674–689 in Thistleton-Dyer W.T., (Ed.) Flora of Tropical Africa 5, Sect. 1, Part 4, London, Lovell & Reeve.

IPNI (continuously updated) The International Plant Names Index. Available from: http://ipni.org/ [accessed Mar. 2018].

IUCN (2012) IUCN red list categories: Version 3.1. Gland, Switzerland and Cambridge, U.K., IUCN Species Survival Commission.

Johnson S.D., Griffiths M.E., Peters C.I., Lawes M.J. (2009) Pollinators, “mustard oil” volatiles, and fruit production in flowers of the dioecious tree Drypetes natalensis (Putranjivaceae). American Journal of Botany 96 (2): 2080–2086. https://doi.org/10.3732/ajb.0800362

Keay, R. W. J. (1958). Euphorbiaceae. In: R. W. J., Keay (ed.) Flora of West Tropical Africa 1: 364–423. London, Crown Agents.

Kirschner J., Cheek M. (2000) New combinations in Tropical African Luzula sect. Luzula (Juncaceae). Kew Bulletin 55: 899–903. https://doi.org/10.2307/4113636

Lachenaud O. (2019) Révision du Genre Psychotria (Rubiaceae) en Afrique Occidentale et Centrale. Opera Botanica Belgica 17: 1–900.

Maisels F.M., Cheek M., Wild C. (2000) Rare plants on Mt Oku summit, Cameroon. Oryx 34: 136–140. https://doi.org/10.1017/s0030605300031057

Mwachala G., Cheek M. (2012) Dracaena mokoko sp. nov. (Dracaenaceae-Ruscaceae/Asparagaceae) a Critically Endangered forest species from Mokoko, Cameroon. Nordic Journal of Botany 30: 389–393. https://doi.org/10.1111/j.1756-1051.2011.01487.x

Nic Lughadha E., Bachman S.P., Govaerts R. (2017) Plant Fates and States: Response to Pimm and Raven. Trends in Ecology & Evolution 32: 887–889. https://doi.org/10.1016/j.tree.2017.09.005

Nic Lughadha E., Govaerts R., Belyaeva I., Black N., Lindon H., Allkin R., Magill R.E., Nicolson, N. (2016) Counting counts: Revised estimates of numbers of accepted species of flowering plants, seed plants, vascular plants and land plants with a review of other recent estimates. Phytotaxa 272: 82–88. https://doi.org/10.11646/phytotaxa.272.1.5

Onana J. (2011) The vascular plants of Cameroon, a taxonomic checklist with IUCN Assessments. Kew, Royal Botanic Gardens.

Onana J., Cheek M. (2011) Red data book of the flowering plants of Cameroon, IUCN global assessments. Kew, Royal Botanic Gardens.

Pax F., Hoffmann K. (1922) Euphorbiaceae - Phyllanthoideae – Phyllantheae – Drypetinae pp. 227–280 in Engler A. (Ed.) Das Planzenreich IV, 147 XV (Heft 81).

Plants of the World Online (continuously updated). Facilitated by the Royal Botanic Gardens, Kew. Published on the Internet; http://www.plantsoftheworldonline.org/ Retrieved 5 Oct. 2019.

Radcliffe-Smith A. (1987) Euphorbiaceae (Part 1) in Polhill R.M. (Ed.) Flora of Tropical East Africa. Rotterdam, Balkema.

Smedmark J.E., Bremer B. (2011). Molecular systematics and incongruent gene trees of Urophylleae (Rubiaceae). Taxon 60: 1397–1406. https://doi.org/10.1002/tax.605015

Soltis D.E, Clayton J.W, Davis C.C. et al. (2007) Monophyly and relationships of the enigmatic family Peridiscaceae. Taxon 56: 65–73.

Sosef M.S.M., Wieringa J.J., Jongkind C.C.H., Achoundong G., Azizet Issembé Y., Bedigian D., Van Den Berg R.G., Breteler F.J., Cheek M., Degreef J. (2006) Checklist of Gabonese Vascular Plants. Scripta Botanica Belgica 35. National Botanic Garden of Belgium.

Tchouto P., Edwards I., Cheek M., Ndam N., Acworth J. (1999) Mount Cameroon Cloud Forest. Pp. 263–277 in African Plants: Biodiversity, Taxonomy, and Uses. Kew, Royal Botanic Gardens.

Thiers B. (continuously updated) Index Herbariorum: A global directory of public herbaria and associated staff. New York Botanical Garden’s Virtual Herbarium. [continuously updated]. Available from: http://sweetgum.nybg.org/ih/ [accessed: March 2018].

Thomas D., Cheek M. (1992) Vegetation and plant species on the south side of Mount Cameroon in the proposed Etinde reserve. Report to Govt. Cameroon from ODA. Kew, Royal Botanic Gardens.

Turland N. J., Wiersema J.H., Barrie F.R., Greuter W., Hawksworth D.L., Herendeen P.S., Knapp S., Kusber W-H., Li D-Z., Marhold K., May T.W., McNeill, J., Monro A.M., Prado J., Price M.J., Smith G.F. (2018) International Code of Nomenclature for algae, fungi, and plants (Shenzhen Code) adopted by the Nineteenth International Botanical Congress Shenzhen, China, July 2017 (eds.). Regnum Vegetabile 159. Glashütten: Koeltz Botanical Books. https://doi.org/10.12705/Code.2018

Willis K.J. (ed.) (2017). State of the World’s Plants 2017. Kew, Royal Botanic Gardens.

